# It’s all in the timing: Delayed feedback in autism may weaken predictive mechanisms during contour integration

**DOI:** 10.1101/2024.01.16.575908

**Authors:** Emily J. Knight, Ted S. Altschuler, Sophie Molholm, Jeremy W. Murphy, Edward G. Freedman, John J. Foxe

## Abstract

Humans rely on predictive mechanisms during visual processing to efficiently resolve incomplete or ambiguous sensory signals. While initial low-level sensory data are conveyed by feedforward connections, feedback connections are believed to shape sensory processing through conveyance of statistical predictions based on prior exposure to stimulus configurations. Individuals with autism spectrum disorder (ASD) show biases in stimulus processing toward parts rather than wholes, suggesting their sensory processing may be less shaped by statistical predictions acquired through prior exposure to global stimulus properties. Investigations of illusory contour (IC) processing in neurotypical (NT) adults have established a well-tested marker of contour integration characterized by a robust modulation of the visually evoked potential (VEP) – the *IC-effect* – that occurs over lateral occipital scalp during the timeframe of the N1 component. Converging evidence strongly supports the notion that this *IC-effect* indexes a signal with significant feedback contributions. Using high-density VEPs, we compared the *IC-effect* in 6–17-year-old children with ASD (n=32) or NT development (n=53). Both groups of children generated an *IC-effect* that was equivalent in amplitude. However, the *IC-effect* notably onset 21ms later in ASD, even though timing of initial VEP afference was identical across groups. This suggests that feedforward information predominated during perceptual processing for 15% longer in ASD compared to NT children. This delay in the feedback dependent *IC-effect*, in the context of known developmental differences between feedforward and feedback fibers, suggests a potential pathophysiological mechanism of visual processing in ASD, whereby ongoing stimulus processing is less shaped by statistical prediction mechanisms.

**SIGNIFICANCE STATEMENT:** Children with autism often present with an atypical visual perceptual style that emphasizes parts or details over the whole. Using electroencephalography (EEG), this study identifies delays in the visual feedback from higher order sensory brain areas to primary sensory regions. Because this type of visual feedback is thought to carry information about prior sensory experiences, individuals with autism may have difficulty efficiently using prior experience and predictions to help make sense of incoming new visual information. This provides empirical neural evidence to support theories of disrupted sensory perception mechanisms in autism.

## INTRODUCTION

Individuals with ASD are notable for an atypical cognitive style, often emphasizing parts rather than wholes (Frith U, 1989). Enhanced perceptual processing of features (Mottron L et al., 2006), weakness in global processing (Happe FG and Booth RD, 2008), or weakened application of prior knowledge to the processing of incoming sensory data (Pellicano E and Burr D, 2012), have been offered as explanations of this characteristic imbalance. Here we report on delayed feedback in the context of unaltered feedforward contributions to early perceptual processing, providing neurophysiologic evidence in support of these theories.

Creating a visual representation of an object confronts at least three major inconveniences: 1) Missing information – the retinal surface is interrupted by the optic nerve and by a network of vasculature (Quigley HA et al., 1990); 2) Ambiguity - one object viewed from different angles projects different shapes upon the retina (Kersten D et al., 2004); 3) Poor conditions – environmental conditions are seldom optimal, such that objects are often seen under poor lighting conditions or are partially occluded by other objects. Both the poverty and ambiguity of incoming signals are thought to be resolved via interactions between sensory representations and prior knowledge (Helmholtz H, 1860/1962).

Although the visual system is characterized as a hierarchy, with lower cortex encoding the most basic features, inputting to successively higher areas which encode ever more complex combinations (Hubel DH and Wiesel TN, 1968), information moves rapidly both up and down the system (Rockland KS and Pandya DN, 1979). Feedforward pathways play a key role in extracting and integrating sensory data (DeYoe EA and Van Essen Dc, 1988), whereas feedback projections are thought to convey statistical predictions based upon prior experience (Rao RP and Ballard DH, 1999). The predictions, in turn, shape the feedforward information via an automatic and rapid iterative process that disambiguates the representation of incoming data (Foxe JJ and Simpson GV, 2002;Kelly SP et al., 2008;Lamme VA et al., 1998;Lee TS et al., 1998;Zipser K et al., 1996).

Feedforward and feedback projections in the visual cortex of non-human primates originate and terminate in different layers of cortex (Rockland KS and Pandya DN, 1979) and crucially, they reach their mature targets over considerably different developmental time courses (Barone P et al., 1995). Prolonged maturation of feedback projections is also seen in humans (Burkhalter A, 1993), establishing a neural basis for the selective vulnerability of the connections they make and their predictive role in visual processing, necessitating exploration of the role of feedback in clinical populations manifesting atypical development.

Toward that end, visual binding paradigms offer an accessible vehicle to probe the integrity of these projections in various populations. Binding of elements in the formation of visual object representations has been associated with feedback in non-human primates (Hupe JM et al., 1998). In humans, delays specific to feedback connections have been associated with visual binding deficits in schizophrenia independently of altered timing in feedforward connections (Kemner C et al., 2009). Contour integration, involving the filling-in between fragments of contours, is one such binding task (Lee TS and Nguyen M, 2001;Murray MM et al., 2002), and this mechanism has been extensively studied using Kanizsa illusory contours (IC) (Kanizsa G, 1976). A modulation of the visual evoked potential (VEP) indexes this process in neurotypical adults and children (Altschuler TS et al., 2012; Murray MM,Wylie GR,Higgins BA,Javitt DC,Schroeder CE and Foxe JJ, 2002;Proverbio AM and Zani A, 2002). This modulation onsets within approximately 90ms of stimulus presentation and peaks at around 150ms in neurotypical adults. This modulation has been termed the *IC-effect,* and is associated with automatic filling-in of object boundaries (Shpaner M et al., 2009).

This *IC-effect* has been localized to the lateral occipital complex (LOC) {Fiebelkorn, 2010 #63;Murray, 2004 #121;Murray, 2006 #56;Murray, 2002 #38}, a group of extrastriate regions that encodes information about coherent objects, independent of the features of which they are comprised (Grill-Spector K et al., 2001). Converging evidence from animal and human work strongly supports a feedback-driven model of contour integration. Studies in non-human primates and mice that have indexed the precise timing of contour integration imply a significant role of feedback connections in this type of processing (Pak A et al., 2020;Zipser K,Lamme VA and Schiller PH, 1996). Likewise, in humans, initial afferent input to visual cortex can be detected using VEPs at between 45-70ms post-stimulation (Foxe JJ and Simpson GV, 2002). Presumably, this initial volley is dominated by representation of local features processed in lower visual areas, feeding forward through the system (Schroeder CE et al., 1998). Rapidly thereafter, higher-order integrative cortices receive initial afferent inputs and begin to convey information about the global scene back to lower levels {Bar, 2006 #217;Chernyshev, 2016 #165;Foxe, 2022 #205;Foxe, 2002 #212;Kelly, 2008 #170;Wokke, 2013 #156}. The timing of the *IC-effect* emerging between 90 and 120ms, suggests that it is largely dominated by feedback processing, and indeed the importance of feedback in contour integration has been confirmed using multiple methodologies including direct disruption of visual feedback with transcranial magnetic stimulation (Pak A,Ryu E,Li C and Chubykin AA, 2020;Wokke ME et al., 2013;Zeng H et al., 2020)

Here we set out to make use of the exquisitely time-sensitive metric of electrophysiological contour integration to investigate these feedback-dominated binding processes across a range of stimulus sizes in a cohort of 6-17-year-old ASD participants. Our central thesis was that these feedback processes would be delayed or disrupted in autism spectrum disorder, offering a potential neural mechanism for the proposed prediction and integration deficits associated with this condition.

## MATERIALS AND METHODS

### Participants

57 NT and 38 ASD individuals aged 6-to-17 years participated. Their sex, age, non-verbal IQ scores, and other pertinent descriptive data are provided in Table 1. Average age and non-verbal IQ scores did not differ between groups. There was a male predominance in the ASD group relative to the control group. Data from 4 neurotypical and 6 ASD participants were excluded either due to poor data quality as evidenced by rejection of greater than 50% of trials or for neuropsychological diagnoses uncovered following recording (in the NT group), resulting in a final cohort of 53 NT and 32 ASD participants. Participants provided informed assent and their parent or guardian gave informed consent. The City College of the City University of New York, Montefiore Medical Center, and Albert Einstein College of Medicine Institutional Review Boards approved all procedures.

**Table 1.**
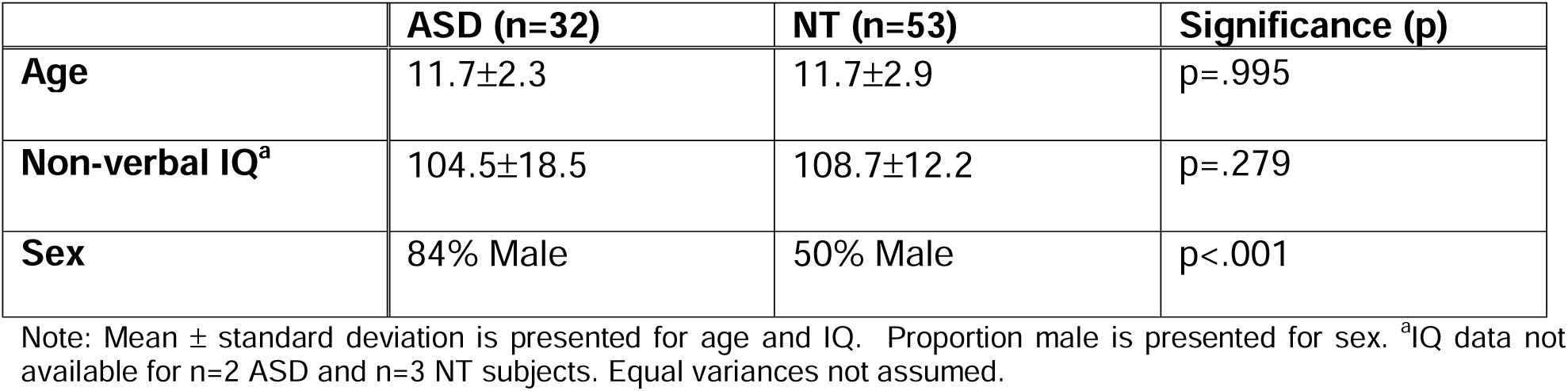
Participant Characteristics.

|  | **ASD (n=32)** | **NT (n=53)** | **Significance (p)** |
| --- | --- | --- | --- |
| **Age** | 11.7±2.3 | 11.7±2.9 | p=.995 |
| **Non-verbal IQ<sup>a</sup>** | 104.5±18.5 | 108.7±12.2 | p=.279 |
| **Sex** | 84% Male | 50% Male | p<.001 |

Note: Mean ± standard deviation is presented for age and IQ. Proportion male is presented for sex. <sup>a</sup>IQ data not available for n=2 ASD and n=3 NT subjects. Equal variances not assumed.

Exclusionary criteria for both groups included history of seizures, head trauma, intellectual disability (full scale IQ <70), schizophrenia, bipolar disorder or psychosis, and history of neurologic disorder or an identified syndromic cause for ASD. Additional exclusionary criteria for neurotypical participants included diagnosis of attention deficit hyperactivity disorder (ADHD), learning disability, other developmental disorder, or history of a developmental disorder in a first-degree relative. Participants were screened for normal or corrected-to-normal vision, hearing, and color vision. Diagnoses of ASD were made on the basis of the Autism Diagnostic Observation Schedule (Lord C et al., 1999) and Autism Diagnostic Interview-R (Lord C et al., 1994) using DSM-IV criteria (assessments collected prior to the 2013 update to the DSM-V). Parents were asked to refrain from giving stimulant medication to their children in the 24 hours preceding participation. Six remaining participants were treated with an antipsychotic and anxiolytic to treat anxiety (1), a mood stabilizer and anti-hypertensive (1) and a norepinephrine reuptake inhibitor (2) to treat ADHD, and a serotonin selective reuptake inhibitor (2) to treat anxiety.

### Stimulus and Task

Participants viewed a version of the Kanizsa illusion, consisting of four black “pacman-like” discs against a gray background {Kanizsa, 1976 #49} (Figure 1). Each disk occupied one of four corners of a square-shaped array. Each had a 90° angle cut out of them - their “mouth.” When the mouths were angled such that their contours were collinear, the gap between the mouths appeared to fill-in, inducing the perception of a square (IC condition). When the mouths were not aligned, no illusion was induced (No-IC condition). In the No-IC condition, three of four inducers are rotated away from the center, the fourth inducer’s location varied randomly and equiprobably. Retinal eccentricity of illusory squares was manipulated randomly and equiprobably within blocks among 3 conditions subtending 4°, 7°, and 10° of visual angle (extent). Inducers for the three extents were 2.1°, 3.8°, and 5.6° diameter respectively, holding support ratio (the proportion of actual to perceived contour extent) constant.

**Figure 1.**
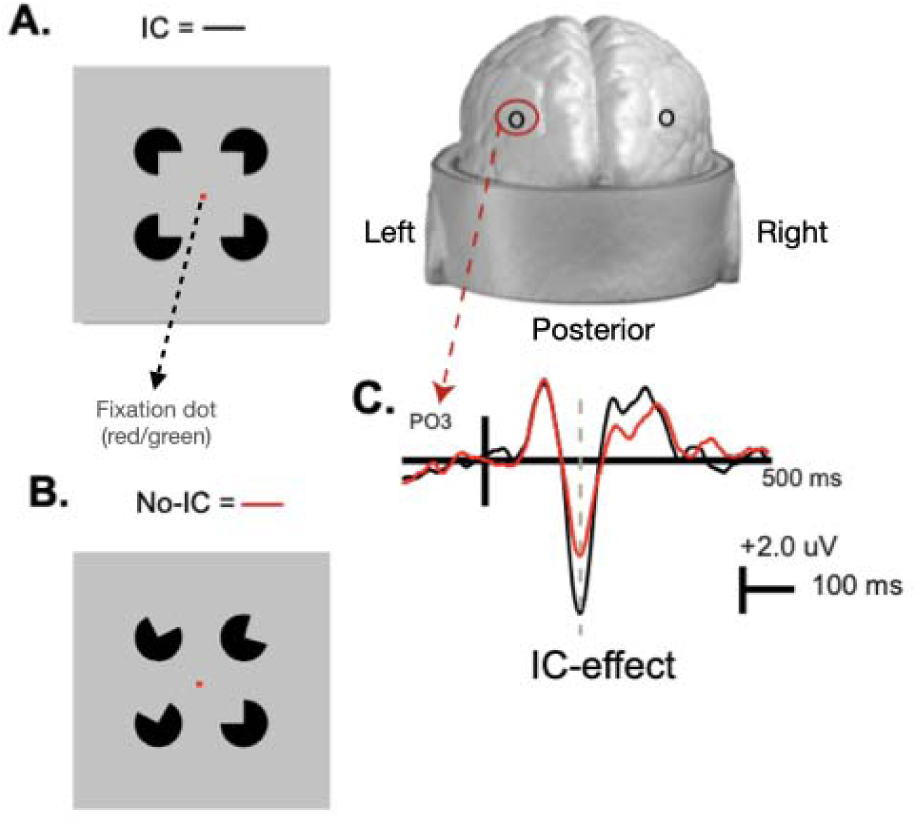
Representation of the A) Illusory Contour (IC; black) and B) Non-contour (No-IC; red) stimuli as well as C) schematic of the *a priori* defined electrodes of interest (PO3 and PO4) and sample visual evoked potential tracings for IC and No-IC stimuli with the *IC-effect* marked by a dashed line.

Participants sat in a dimly lit, sound-attenuated electrically shielded double-walled booth (Industrial Acoustics Company, Bronx, NY), 60 cm from an LCD monitor with 1280 x 1024-pixel resolution or 75 cm from a monitor with 1680 x 1050 pixel resolution. Stimulus durations were 500ms with an 800–1400ms onset asynchrony, varied on a square wave distribution. Ten to fifteen 3-minute blocks were presented, with breaks as needed, until sufficient (>100 trials per condition) had been collected. Explicit attention to ICs is not required to elicit electrophysiological indices of IC processing in NT adults (Murray MM,Wylie GR,Higgins BA,Javitt DC,Schroeder CE and Foxe JJ, 2002) or children (Altschuler TS et al., 2014) when stimuli are centrally presented (i.e. foveated). Task instructions made no mention of the specific stimuli or the illusion. Instead, participants attended to a color-detection task involving the central fixation dot. Every 1-10 seconds, the dot changed from red to green for 160ms on a random time-course uncorrelated with IC presentation. As the colors were chosen for an isoluminant plane of the DKL color-space (Derrington AM et al., 1984) the change was imperceptible without foveating. Participants were asked to click a mouse button for each color-change. All groups performed well above chance, but participants with ASD performed less well (Mean Accuracy: ASD 84.2 ± 12.7%, NT 90 ± 12.5%, *t*_(79*)_ = 2.334; p =.022, Cohen’s d=.531, *four NT participants for whom behavioral data was not stored were excluded from this analysis). 6-9-year-old participants were observed to ensure fixation.

Following administration of the main VEP experiment, a debriefing questionnaire assessed IC perception. When shown square-inducing stimuli like that used in the experiment, 100% of included participants identified the IC-condition as the “square.”

### Data Acquisition and Processing

Continuous EEG was recorded via a Biosemi ActiveTwo system from a 70-electrode montage, digitized at 512 Hz and referenced to the Common Mode Sense (CMS) and Driven Right Leg (DRL). EEG data were processed and analyzed offline using custom scripts that included functions from the EEGLAB (Delorme A and Makeig S, 2004) and ERPLAB Toolboxes (Lopez-Calderon J and Luck SJ, 2014) for MATLAB (the MathWorks, Natick, MA, USA). Data were band-pass filtered using an IIR Butterworth filter between 0.1 and 50 Hz implemented in ERPLAB. Bad channels were manually and automatically detected and interpolated using EEGLAB spherical interpolation. Data were re-referenced to a frontal electrode (Fpz in the 10-20 system convention) and then divided into epochs starting 100ms before the presentation of each IC/No-IC stimulus and extending to 500ms post-stimulus onset. Trials containing severe movement artifacts or particularly noisy events were rejected if voltages exceeded ±125μV. Trials were then averaged to obtain grand average waveforms for 4°, 7°, and 10° IC and No-IC stimulus presentations for each subject. Median number of interpolated channels and accepted trials per condition for each group is shown in Table 2. Analyses were guided by previous IC work (Altschuler TS,Molholm S,Butler JS,Mercier MR,Brandwein AB and Foxe JJ, 2014) which identified ERP effects sensitive to the difference between IC conditions during time windows associated with the visual N1.

**Table 2.** Median number of channels interpolated and number of trials accepted per group.

|  | **ASD (n=32)** | **NT (n=53)** | **Significance (p)** |
| --- | --- | --- | --- |
| **Channels Interpolated** | 4 (1-11) | 4 (1-11) | p=.737 |
| **Trials Accepted** | 1375 (777-1689) | 1419 (839-1917) | p=.128 |

Note: Median (Range) are reported. Number of trials accepted includes both IC and No-IC conditions at all stimulus sizes. Statistical significance evaluated via Mann Whitney U due to skewed distributions.

#### Primary analysis

##### Group Comparison of IC-effect amplitudes

Statistical analyses were implemented in SPSS (IBM Corp. Released 2020. IBM SPSS Statistics for MacOS, Version 27.0. Armonk, NY: IBM Corp). To examine contour integration for Kanizsa figures of varying support ratios in ASD and NT participants, while limiting type-II errors, the initial analysis was restricted both spatially and temporally. We focused on electrodes over lateral occipital scalp sites (PO3 and PO4) where prior literature has suggested the strongest *IC-effect* responses are observed (Altschuler TS,Molholm S,Russo NN,Snyder AC,Brandwein AB,Blanco D and Foxe JJ, 2012;Foxe JJ et al., 2005). Data were first collapsed across both sensors of interest and diagnostic groups. The time window for the *IC-effect* and was initially broadly defined based on component latency windows described in a previous study mapping the spatiotemporal dynamics of IC processing in NT children (Altschuler TS,Molholm S,Butler JS,Mercier MR,Brandwein AB and Foxe JJ, 2014) and then further refined within these general component time windows by the grand-averaged waveforms collapsed across both groups, inclusive of IC and No-IC stimuli (i.e. without regard for or bias from the dependent measures of interest). Mean amplitudes were then computed for each subject over a 10ms time window (184 to 194ms) for the *IC-effect*. Finally, we implemented a mixed model analysis of variance (ANOVA) with a between-subjects factor of group (NT, ASD) and within-subject factors of stimulus size (4°,7°, 10°) and hemisphere (Left-PO3, right PO4) to compare the *IC-effect* mean amplitude between groups.

#### Secondary Analyses

Given that hemispheric lateralization and stimulus size did not appear to differentially modulate the *IC-effect* between groups, data were collapsed across all conditions for these exploratory secondary analyses.

##### Group Comparison IC-effect onset latencies

We noted when viewing the data that although both groups generated an *IC-effect* that was robust in amplitude (contrary to our initial prediction), the *timing* of this processing appeared to differ between groups. As a result, we conducted an exploratory analysis to assess the magnitude of these latency differences. Because difference waves necessarily have a lower signal-to-noise ratio, we used a jackknife-based method to estimate onset latencies of the *IC-effect*. This method has been shown to outperform methods based on the selection of onset latencies at the single participant level (Miller J et al., 1998;Ulrich R and Miller J, 2001). It proceeds as follows: for all *n* subjects in a given group, 1 subject is omitted, and the average computed over the remaining *n - 1* subjects. *n* averages are computed, each subtracting 1 subject’s data. For each of these *n* jackknife waveforms, an onset latency was computed. Onset latency was defined as the point between the predefined window of 50-250ms at which the voltage reached 50% of the minimum voltage, a well-accepted estimate of onset previously used for difference-wave measures (Luck SJ et al., 2009). Since the jackknife waveforms are digitally sampled, and thus discrete, we rarely possessed a sampled latency value that corresponded precisely with the 50% criterion. As such, we linearly interpolated between the nearest two latency values (above and below) the precise 50% voltage value. All jackknife latency measurements were conducted using ERPLAB built-in functions (Lopez-Calderon J and Luck SJ, 2014). Ulrich and Miller (Ulrich R and Miller J, 2001) rigorously demonstrated that the jackknife technique artificially reduces the error variances in the ANOVA. Therefore, prior to conducting the statistical analysis we used the method outlined by Smulders (Smulders FT, 2010) to extract individual latencies from the jackknife average waveforms, allowing for application of traditional parametric statistical testing. These extracted latencies were then compared between groups using a Student’s t test.

##### Group Comparison VEP onset latencies

To confirm that any changes in feedback-associated processes during the N1 latency were not due to differences at the onset of cortical visual processing, we compared the onset latency of the P1 for each group at each of the pre-defined electrode sites (left-PO3, right-PO4). For each jackknife waveform, we calculated the average onset latency for the P1 evoked by IC/No-IC stimuli. P1 onset latency was defined as the point between the predefined window of 0-180ms at which the voltage reached 50% of the maximum voltage. As above, when a sampled latency value did not correspond with the 50% criterion, we linearly interpolated between the nearest two latency values (above and below) the precise 50% voltage value. Following the same methodology as above, we extracted individual latencies from the jackknife average waveforms and compared these between groups using a Student’s t test.

##### Sex modulation of IC-effect

We also noted a significant difference in sex distribution between the two groups with a male predominance in the ASD group (see Table 1). Therefore, to evaluate whether sex influences the amplitude or latency of the *IC-effect* in children, Student’s t-tests were performed to compare the amplitude and onset latency (calculated via the methodologies outline above) of the *IC-effect* between males and females within the NT group only.

## RESULTS

### IC-effect amplitude

Grand-average VEPs to the IC and No-IC stimulus configurations at the *a-priori* defined electrodes of interest (PO3 and PO4), averaged across all stimulus sizes, are depicted for each group in Figure 2A-B, and the results of the primary analysis are summarized in Table 3. As evident in Figure 2, the IC stimuli evoked stronger negative responses than No-IC stimuli in the N1 time window in both groups, the so-called *IC-effect*. For the *IC-effect* time windows, there were no overall differences in the magnitude of the IC effect between ASD and NT groups (F(1,82)=3.038, p=.089, η_p_^2^=.034) and no group-related interactions. There was no significant hemispheric lateralization of the *IC-effect* (F(1,83)=2.564, p=.113, η_p_^2^=.030). However, the magnitude of the *IC-effect* was modulated by size with larger stimulus size generating a stronger *IC-effect* (F(2,166)=4.084, p=.019, η_p_^2^=.047) in both groups. To allow for visualization of the impact of stimulus size, topographic maps depicting the differences in evoked response to IC minus No-IC stimuli for each of the three stimulus sizes (4°,7° and 10°), along with associated grand-average VEPs to the IC and No-IC stimulus configurations at the a-priori defined electrodes of interest (PO3 and PO4), are presented in Figure 3.

**Figure 2.**
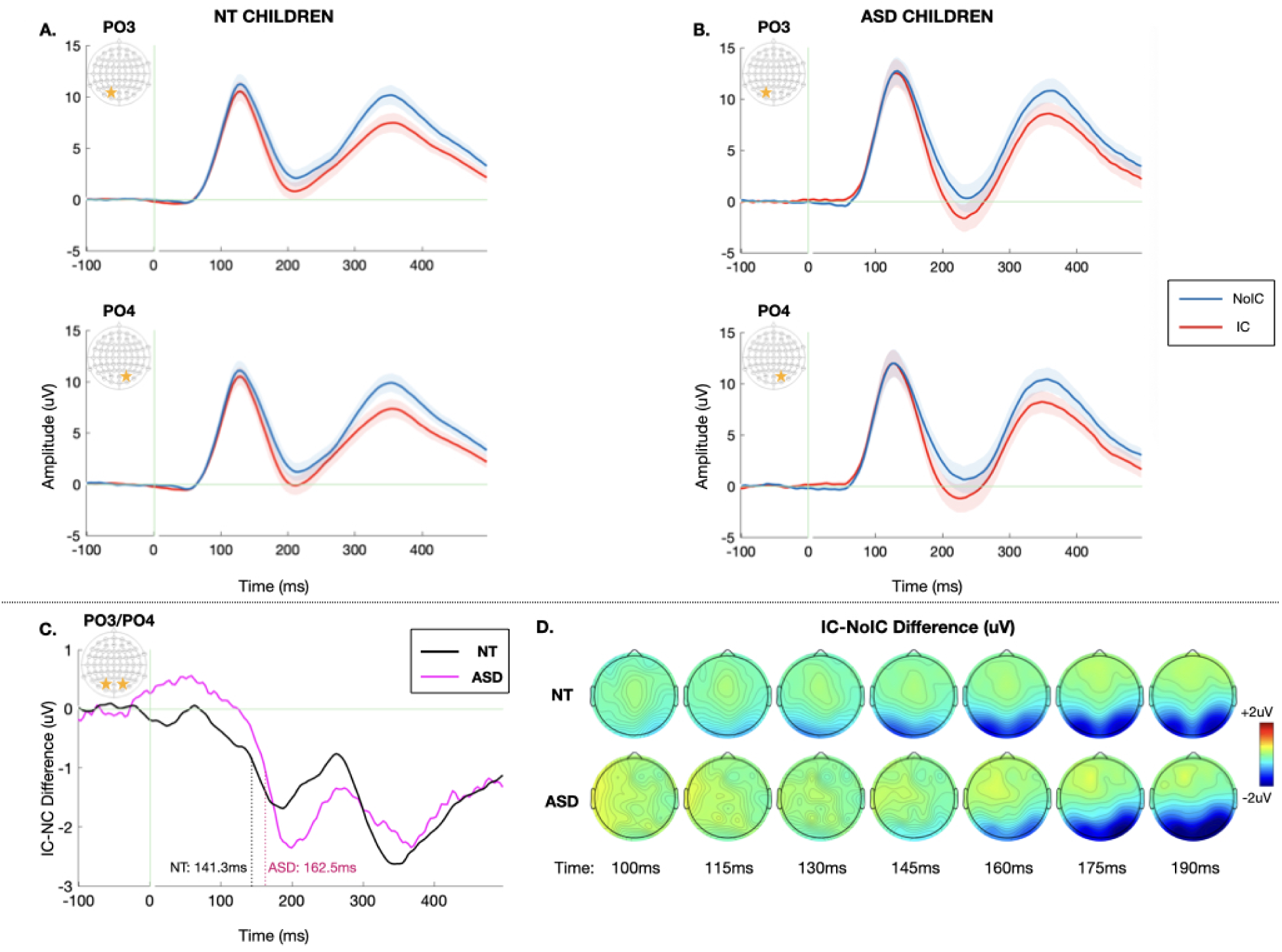
Grand average visual potentials evoked by non-contour/No-IC (blue) and illusory contour/IC (red) stimuli over the left (electrode PO3) and right (electrode PO4) lateral occipital regions for the A) NT B) ASD groups. Data are averaged across all stimulus sizes. Shaded regions depict standard error of the mean (SEM). Green vertical line marks stimulus onset at t=0. C) *IC-effect* difference wave (IC minus No-IC) averaged across PO3 and PO4. *IC-effect* onset derived from jackknifed measures for each group (50% of the minimum difference) is marked by dotted lines for the ASD (pink) and NT (black) groups. D) Topographic maps depicting the *IC-effect,* or the difference in amplitude evoked by IC – No-IC stimuli between 100-190ms for the NT (top) and ASD (bottom) groups.

**Figure 3.**
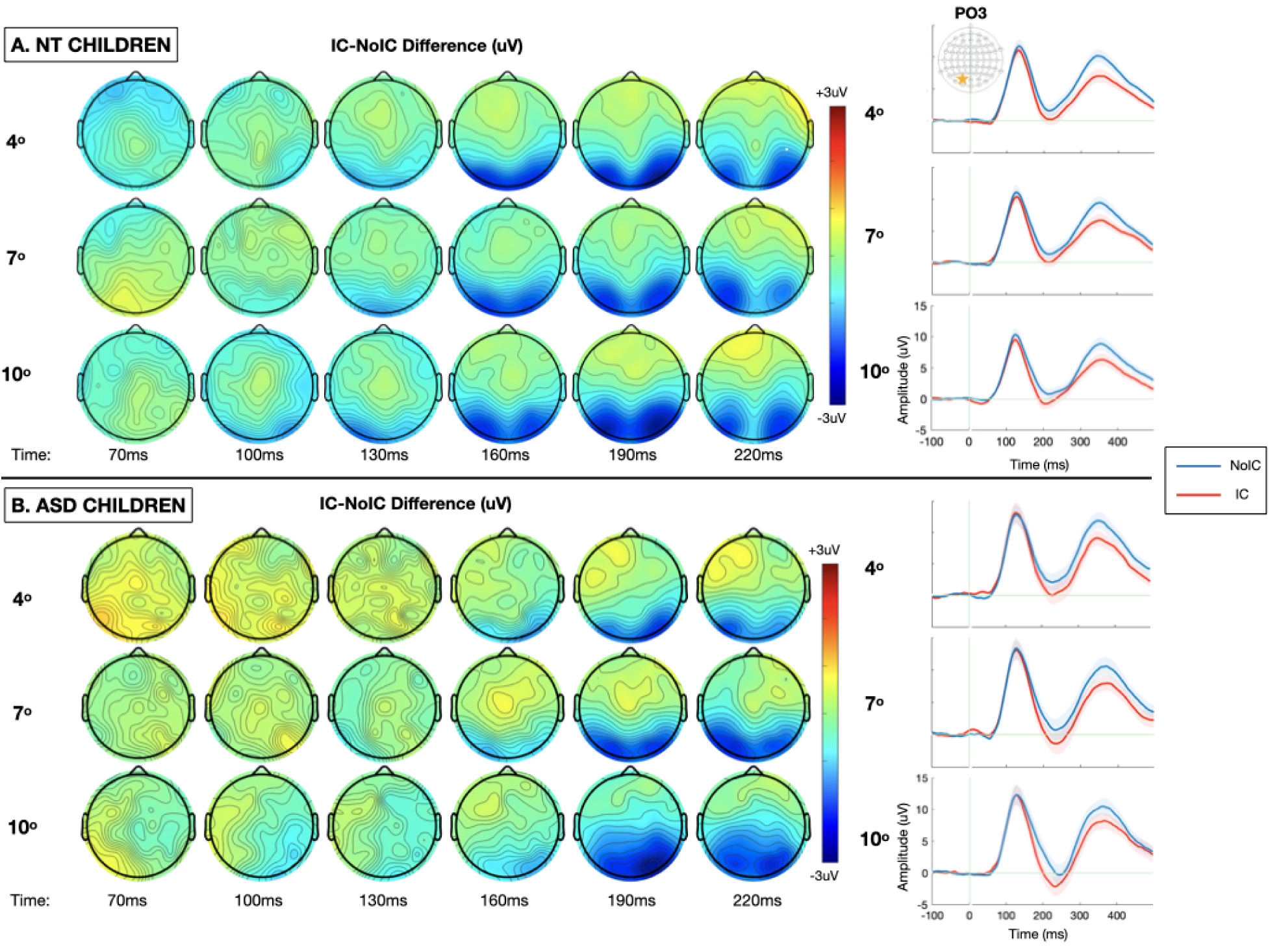
Topographic maps depicting the *IC-effect,* or the difference in amplitude evoked by IC – No-IC stimuli, between 70-220ms for each of the three stimulus sizes (4, 7, and 10° visual angle), alongside the corresponding grand average visual potentials evoked by non-contour/No-IC (blue) and illusory contour/IC (red) stimuli over the left lateral occipital region (electrode PO3) for the (A) NT (B) ASD groups.

**Table 3.** Primary Analysis: IC-Effect Amplitude.

|  | *F* | *IC-effect* | *p-value* |
| --- | --- | --- | --- |
| **Main Effects** |  |  |  |
| Group | 2.960 |  | .089 |
| Hemisphere | 2.564 |  | .113 |
| Stimulus Size (4°, 7°, 10°) | 4.084 |  | **.019\*** |
| **Within Subjects Interactions** |  |  |  |
| Hemisphere x Stimulus Size | 1.275 |  | .282 |
| **Within Subject x Between Subject Interactions** |  |  |  |
| Hemisphere x Group | .500 |  | .481 |
| Stimulus Size x Group | 1.260 |  | .286 |
| Hemisphere x Stimulus Size x Group | .577 |  | .574 |

Note: \* $p<.05$ ; $n_{NT}=53$ , $n_{ASD}=32$

To highlight the timing of contour integration, the difference in evoked response to IC minus No-IC stimuli at the two electrodes of interest (PO3 and PO4) for the ASD and NT groups are depicted in Figure 2C, alongside topographic maps allowing for visualization of effects across the entire array (Figure 2D). Onset latency of the *IC-effect* was estimated at 162.5 ± 27.1ms for participants with ASD and 141.3 ± 54.5ms for NT participants. This 21ms delay yielded a main effect of diagnosis (t_83_=2.383*, p=.020, Cohen’s d= .458, *equal variances not assumed due to significant Levene’s Test for equality of variances) (Figure 2C). In contrast to the latency of the *IC-effect,* onset latency of the P1 was estimated at 94.3± 14.3ms for participants with ASD and 96.3± 32.4ms for NT participants, and thus clearly not significantly different between groups (t_83_=-3.86*, p=.747, Cohen’s d= -.086, *equal variances not assumed due to significant Levene’s Test for equality of variances). Thus, NT participant’s IC processing was feedforward-dominated for approximately 45ms, whereas participants with ASDs was feedforward-dominated for 68ms. *IC-effect modulation by sex* When comparing the *IC-effect* between males and females within the NT group, there was no effect of sex on amplitude (t_50_ = -.498, p=.621, Cohen’s d=-.138) or latency (t_50_ = -.686, p=.496, Cohen’s d=-.190).

## DISCUSSION

To investigate feedback contributions to visual processing in ASD, we compared a well-tested metric of automatic contour completion in 6–17-year-olds with ASD and their neurotypical counterparts. The *IC-effect* generated was equivalent in amplitude between ASD and NT groups. However, the *IC-effect* onset 21ms later in individuals with ASD despite simultaneous onset of visual cortical activity across the two groups. This pattern of results is suggestive of delayed visual feedback processing in ASD.

Feedforward and feedback connections between human visual cortical areas V1 and V2 seem to develop from segregated populations of neurons and follow different developmental growth patterns (Burkhalter A, 1993). While feedforward axons grow toward their target cortical layers precisely, reaching them by approximately 4 months of age, feedback fibers grow past their targets, sending out multiple buds from the axon and have still not reached their targets by this time. Synaptic proliferation (Huttenlocher PR and de Courten C, 1987) and growth of dendritic spines (Michel AE and Garey LJ, 1984) increase in humans over the first five months of life, suggesting that space for feedback inputs may not be available until later in infancy (Burkhalter A, 1993;Rabinowicz T, 1986). Brain overgrowth prior to three years of age is a consistently replicated finding in a substantial subset of children with ASD (Hazlett HC et al., 2011;Yankowitz LD et al., 2020). It has been connected to increased neuron count (Courchesne E et al., 2011) and density (Hutsler JJ and Zhang H, 2010). One possibility is that initial overexuberant feedforward connections cause feedback fibers to encounter greater obstacles to reaching their intended targets. Protracted development of feedback circuitry would likely alter the predictive function of feedback circuitry in sensory processing (Rao RP and Ballard DH, 1999), particularly if it is dependent on exposure gained through the earlier maturing feedforward circuitry (Berezovskii VK et al., 2011).

Typically, ambiguous sensory inputs are shaped by statistical predictions about configuration acquired through prior exposure (Lee TS and Mumford D, 2003). Delayed onset of feedback dominated activity in ASD suggests that perceptual representations may remain less shaped by such internal input. Local features influence initial perceptual processing for a longer time – in this case for ∼15% longer – indicating reduced influence of priors on processing of local stimulus elements. Notably, similar patterns of delayed latency but normal amplitude visual evoked potentials have been implicated in global processing for subjects with schizophrenia, a neurologic condition with shared genetic susceptibility and symptom overlap with ASD (Kemner C,Foxe JJ,Tankink JE,Kahn RS and Lamme VA, 2009).

Reduced predictive feedback does not imply only negative outcomes – that would depend on the perceptual task to which they contribute. The ambiguity of incoming features may simply be resolved later; alternatively, visual processing may adapt to more feedforward-weighted input. Such input may explain why individuals with ASD excel in tasks like the copying of geometrically impossible figures (Mottron L et al., 1999), where delayed feedback may mean that copying is less influenced by prior knowledge. Assuming this pattern predominated over childhood, this could foster development of experience-dependent visual processes that rely less on predictive mechanisms overall. In such a system it may be adaptive to place greater reliance on sensory details than on information about wholes – a characteristic of the ASD phenotype; however, additional work is clearly indicated to understand how the maturational trajectory of the visual feedback mechanisms relates to specific phenotypic characteristics in ASD. Models of predictive processing posit that in situations where sensory input is ambiguous, multiple interpretations may be actively represented in lower level cortex until feedback suppresses those determined to be less likely based on prior knowledge (Lee TS and Mumford D, 2003;Pollen DA, 1999). It is possible to extrapolate that if predictive mechanisms are delayed or weakened, the sensory processing systems of individuals with ASD may be overloaded with an abundance of potential representations.

Interestingly, this work differs from prior work conducted by our group involving children in the same age range, which found a decreased amplitude of the *IC-effect* among autistic children, without clear latency differences (Knight EJ et al., 2023). While both findings point to altered feedback processing during contour integration in ASD, the different manifestations may be due to one of two categories of factors-subject heterogeneity or paradigmatic differences. Examining these areas of agreement and discrepancy are of great interest to help enrich the understanding of the factors influencing contour integration in ASD and the replicability of these findings across studies. Exploration of this type of heterogeneity represents an area of increasing emphasis for electrophysiologic research in ASD (Webb SJ et al., 2015). When comparing these two highly similar studies (Knight EJ,Freedman EG,Myers EJ,Berruti AS,Oakes LA,Cao CZ,Molholm S and Foxe JJ, 2023) on a number of subject factors that might influence the results, including age, sex, and IQ, the subject populations did not appear to differ on any of these factors. Consistent across both studies along with much prior visual perception work, VEP component amplitudes were greater for younger participants in both groups without age-related differences in the developmental trajectory of contour integration mechanisms. The overall larger amplitudes in younger participants may simply be attributable to anatomic features such as skull thickness (Adeloye A et al., 1975).

One possible phenotypic confounder is that participants in this study were diagnostically characterized prior to the implementation of the DSM-5 and are therefore classified under the older DSM-4 criteria. While there is substantial overlap between diagnoses made under the DSM-5 and DSM-4 criteria, those diagnosed with pervasive developmental disorder, not otherwise specified (PDD-NOS) under the DSM-4 have higher rates of loss of autism diagnosis under the newer DSM-5 criteria (Daniels AM et al., 2011). When removing this less stable diagnostic category (n=5) from the results, the pattern of findings did not appear substantially different (See Figure 4 for an alternative visualization of the data excluding those participants with a PDD-NOS diagnosis). Nevertheless, we cannot completely rule out that subject populations between this and the prior similar study (Knight EJ,Freedman EG,Myers EJ,Berruti AS,Oakes LA,Cao CZ,Molholm S and Foxe JJ, 2023) differed on another unmeasured factor, as autism is a highly complex and heterogeneous condition. Regarding paradigmatic differences, spatial attention is potentially an interesting factor {Martinez, 2007 #219;Martinez, 2006 #218;Senkowski, 2005 #182}. While both paradigms involved automatic processing of illusory contour stimuli, the prior study varied the stimulus presentation location which may have resulted in differences in covert spatial attentional orienting. By contrast, in the present study spatial attention demands were limited with all stimuli being presented surrounding central fixation. Indeed, differences in spatial orienting are common (Ciesielski KT et al., 1990;Kawakubo Y et al., 2007;Keehn B et al., 2013;Landry R and Bryson SE, 2004;Sacrey LA et al., 2014) in autism and are likely an underrecognized factor in many studies of sensory perception in ASD.

**Figure 4.**
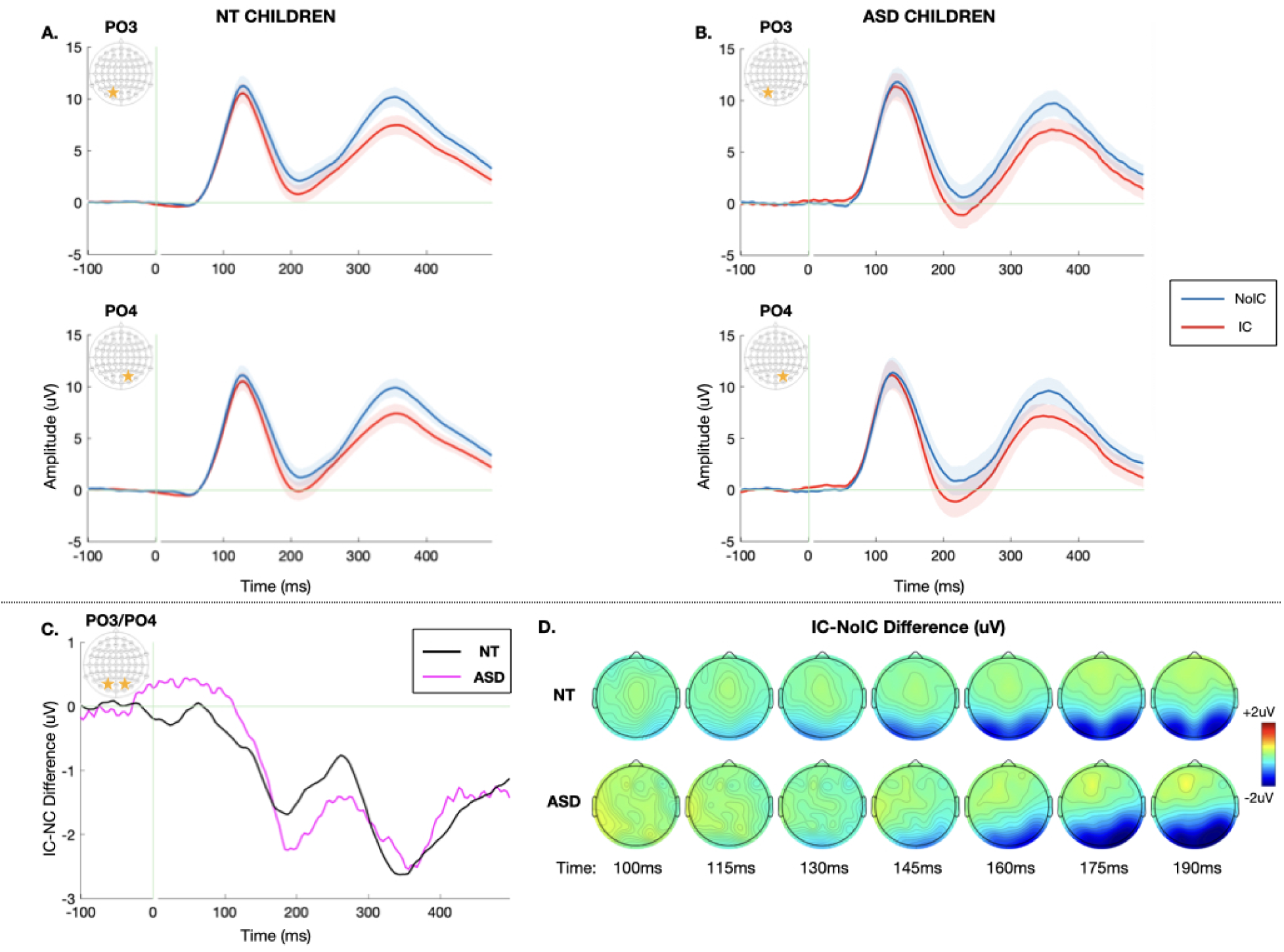
Alternate representation of the data, excluding those participants (n=5) with a diagnosis of pervasive developmental disorder, not otherwise specified (PDD-NOS). Grand average visual potentials evoked by non-contour/No-IC (blue) and illusory contour/IC (red) stimuli over the left (electrode PO3) and right (electrode PO4) lateral occipital regions for the A) NT B) ASD groups. Data are averaged across all stimulus sizes. Shaded regions depict standard error of the mean (SEM). Green vertical line marks stimulus onset at t=0. C *IC-effect* onset derived from jackknifed measures for each group (50% of the minimum difference) is marked by dotted lines for the ASD (pink) and NT (black) groups. D) Topographic maps depicting the *IC-effect,* or the difference in amplitude evoked by IC – No-IC stimuli between 100-190ms for the NT (top) and ASD (bottom) groups.

Here, the main stimulus manipulation was stimulus size ranging from 4° to 10° visual angle, while holding the support ratio constant. Of note, the 4°stimulus is the same size as the IC stimuli presented by (Knight EJ,Freedman EG,Myers EJ,Berruti AS,Oakes LA,Cao CZ,Molholm S and Foxe JJ, 2023). With increasing stimulus size, we observed a stepwise increase in amplitude of the VEP components, including the N1, evoked by IC and No-IC stimuli. This is consistent with prior work suggesting increased recruitment of neurons when there is a wider central stimulus field resulting in increased VEP amplitude (Yadav NK et al., 2012). However, the strength of the *IC-effect* as indexed by the difference in N1 amplitude evoked by IC versus No-IC stimuli remained constant, suggesting that the contour integration response is size invariant over this range of visual angles, again consistent with prior work {Altschuler, 2012 #39;Mendola, 1999 #128}. While one might expect larger stimuli to be biased toward involvement of higher order areas, associated with more pronounced visual feedback and augmented *IC-effect*, the findings are not consistent with that. However, all stimuli here did extend across the vertical meridian, requiring integration across anatomically separate visual processing regions. Our research group has previously demonstrated that presenting illusory contours laterally to central fixation, such that they no longer straddle the vertical meridian, does shift the bias toward more feedforward processing {Murray, 2002 #38;Senkowski, 2005 #182}. Most notably for the purposes of this study, the degree of size-related VEP modulation was equivalent between diagnostic groups, indicating that the ASD participants were not impaired in their ability to integrate the contours over greater visual distances.

Taken together with prior literature, this electrophysiologic work points to deficits in feedback-supported contour integration in ASD (Knight EJ,Freedman EG,Myers EJ,Berruti AS,Oakes LA,Cao CZ,Molholm S and Foxe JJ, 2023;Stroganova TA et al., 2007) that are evident despite conflicting results behaviorally on whether IC processing deficits are present in autism. Some studies have found no difference between ASD and NT on behavioral measures of contour integration (Gowen E et al., 2020;Hadad BS et al., 2019;Milne E and Scope A, 2008), while other studies have described reduced accuracy and longer reaction times in these tasks (Nayar K et al., 2017;Soroor G et al., 2022). This discrepancy between behavioral and electrophysiologic investigation suggests that individuals with ASD may accomplish equivalent IC perception, albeit via different mechanisms than their NT counterparts. Indeed, in a debriefing questionnaire, participants overwhelmingly accurately identified an illusory shape indicating that they were able to see the illusion despite the clear delays in visual feedback. Notably, all paradigms that rely on behavioral identification of illusory contour presence or absence necessarily force explicit attention to contour integration whereas electrophysiologic paradigms allow for examination of automatic contour integration processing while attention is directed elsewhere. It remains a possibility that neural mechanisms of contour integration are augmented when participants are explicitly attending to illusory contour presence/absence, a phenomenon that has been described in other types of global visual perception in autism (Knight EJ et al., 2022). Additional studies to compare across developmental disability populations such as ADHD and to directly assess the role of attention by comparing contour integration in attended vs. passive processing would be highly interesting.

While this study contributes to our understanding of the timing of contour integration processing in ASD, there remain limitations to the scope of the study that are important to highlight. For one, participants in the study represent only a portion of the extensive phenotypic variability that characterizes the autism spectrum. Participants were actively engaged in the color change discrimination task while participating in this study. As a result, included children needed to have the cognitive and verbal ability to understand and comply with these task instructions. Thus, results should be generalized with caution to children with so-called “profound” autism. Future adaptation of fully passive paradigms may permit the inclusion of this understudied population. Additionally, there is substantial overlap between ASD and ADHD. As a result, we are unable to determine whether the reduced contour integration noted in this study is specific to ASD or characteristic of ADHD or other developmental diagnoses as well.

### Conclusion

Here, we demonstrate a 21ms delay in the onset of feedback-dominated visual processing suggesting a mechanism of weakened influence of prior experience in individuals with ASD, which may disrupt the predictive apparatus relied on for rapid, automatic grouping of incoming sensory information.

## Abbreviations List

IC: Illusory Contour
NT: Neurotypical
VEP: Visual Evoked Potential
ASD: Autism Spectrum Disorder
No-IC: Non-Contour
EEG: Electroencephalography
LOC: Lateral Occipital Cortex
ADHD: Attention Deficit Hyperactivity Disorder ANOVA Analysis of Variance
PDD-NOS: Pervasive developmental disorder, not otherwise specified

## Acknowledgements

The authors express their gratitude to Drs Juliana Bates, Hilary Gomes, John S. Butler and Adam Snyder for their expertise and Ms. Sarah Ruberman and Mr. Frantzy Acluche for their hands-on assistance. In addition, without the gracious donation of many hours from the children and their families who participated in this research, it could never have been conducted.

## Ethics and Consent

The City College of the City University of New York, Montefiore Medical Center, and Albert Einstein College of Medicine Institutional Review Boards approved all the experimental procedures. Each child participant provided informed assent and their parent or guardian provided written informed consent in accordance with the tenets laid out in the Declaration of Helsinki.

## Authors’ Contributions

E.J.K. analyzed the data, created the figures, and wrote the first compiled draft of the submitted manuscript. T.S.A developed the experimental paradigm software, collected the data, and wrote an earlier draft of the manuscript. J.W.M developed the procedures for the jackknife latency analysis. S.M and J.J.F. designed the research. E.G.F and J.J.F. supervised the preparation of the final manuscript. E.J.K, T.S.A., S.M, J.W.M, E.G.F, and J.J.F provided input into the editing and final approval of the manuscript.

